# SAMHD1 regulates human papillomavirus 16 induced cell proliferation and viral replication during differentiation of oral keratinocytes

**DOI:** 10.1101/522250

**Authors:** Claire D. James, Apurva T. Prabhakar, Michael R. Evans, Xu Wang, Molly L Bristol, Kun Zhang, Renfeng Li, Iain M. Morgan

## Abstract

Human papillomaviruses induce a host of anogenital cancers, and also oropharyngeal cancer (HPV+OPC); HPV16 is causative in around 90% of HPV+OPC. Using TERT immortalized “normal” oral keratinocytes (NOKs) we have identified significant host gene reprogramming by HPV16 (NOKs+HPV16), and demonstrated that NOKs+HPV16 support late stages of the viral life cycle. Expression of the cellular dNTPase and homologous recombination factor SAMHD1 is transcriptionally regulated by HPV16 in NOKs, and here we demonstrate that E6 and E7 regulate expression of SAMHD1 at the transcriptional and post-transcriptional levels. CRISPR/Cas9 removal of SAMHD1 from NOKs and NOKs+HPV16 demonstrate that SAMHD1 controls cell proliferation of NOKs only in the presence of HPV16; deletion of SAMHD1 promotes hyper-proliferation of NOKs+HPV16 cells in organotypic raft cultures but has no effect on NOKs. Viral replication is also elevated in the absence of SAMHD1. This new system has allowed us to identify a specific interaction between SAMHD1 and HPV16 that regulates host cell proliferation and viral replication; such studies are problematic in non-immortalized primary oral keratinocytes due to their limited lifespan. To confirm the relevance of our results we repeated the analysis with human tonsil keratinocytes immortalized by HPV16 (HTK16) and observe the same hyper-proliferative phenotype following CRISPR/Cas9 editing of SAMHD1. Identical results were obtained with three independent CRISPR/Cas9 guide RNAs. The isogenic pairing of NOKs with NOKs+HPV16, combined with HTK16, presents a unique system to identify host genes whose products functionally interact with HPV16 to regulate host cellular growth in oral keratinocytes.

**Importance:** Head and neck cancer is the sixth most common cancer worldwide. The incidence of HPV+OPC has been rising steadily since the 1970s and has recently reached epidemic proportions, according to the WHO. Upwards of 70% of the 600,000 new OPC cases per year are HPV positive, with high-risk type 16 present in 90% of those incidences. A better understanding of the viral life cycle will facilitate the development of novel therapeutics to combat this ongoing epidemic, as well as other HPV positive cancers. Here we present a unique oral keratinocyte model to identify host proteins that specifically interact with HPV16. Using this system, we report that a cellular gene, SAMHD1, is regulated by HPV16 at the RNA and protein level in oral keratinocytes. Elimination of SAMHD1 from these cells using CRISPR/Cas9 editing promotes enhanced cellular proliferation by HPV16 in oral keratinocytes and elevated viral replication, but not in keratinocytes that do not have HPV16. Our study demonstrates a specific intricate interplay between HPV16 and SAMHD1 during the viral life cycle and establishes a unique model system to assist exploring host factors critical for HPV pathogenesis.

## Introduction

High-risk human papillomaviruses (HPVs) are the etiologic agents of a number of cancers, including anogenital and oropharyngeal carcinomas (OPC) (1). Upwards of 70% of new OPC cases per year are HPV positive, with high-risk type 16 responsible for 90% of these HPV positive cases (2). There have been extensive studies investigating the HPV16 life cycle in ano-genital keratinoctyes (for example, (3–6)). Such studies have been crucial in enhancing our understanding of the viral life cycle and identifing novel therapeutic targets that could be targeted for alleviation of HPV induced cancers and/or disruption of the viral life cycle. While HPV+ cervical and oropharyngeal cancers have the same etiological agent, there are distinct differences between the diseases. For example, the majority of HPV+OPC retain episomal genomes replicating in an E1-E2 dependent manner whereas in cervical cancer most often the viral genome is integrated into that of the host (7, 8). Such differences, and the ongoing epidemic of HPV+OPC, prompted us to develop and characterize an oral keratinocyte model supporting late stages of the HPV16 life cycle in organotypic raft cultures (9). This was done by introducing the HPV16 genome into TERT immortalized “normal” oral keratinocytes (NOKs, generated from a 28 year old male (10)) and carrying out organotypic raft cultures followed by confirmation of late stages of the viral life cycle including E1^E4 and E2 expression as well as amplification of the HPV16 genome in the differentiated epithelial cells. We used this model to determine host reprogramming induced by HPV16 in oral keratinocytes and a large number of innate immune genes were shown to be downregulated, as others have demonstrated before (9). One of the innate immune genes predicted to be downregulated was SAMHD1 (Sterile alpha motif and histidine-aspartic domain HD containing protein 1). To our knowledge this is the first time downregulation of SAMHD1 expression by HPV16 has been reported.

SAMHD1 is a triphosphohydrolase (dNTPase) enzyme, which regulates intracellular levels of dNTPs and acts as an intrinsic immune response factor (11, 12). To function as a dNTPase, the protein forms a homo-tetramer, which is destabilized by phosphorylation. Each individual protein is comprised of two domains: a sterile alpha motif (SAM), which mediates protein-protein contacts often with other SAM domains, and a dGTP-regulated dNTP hydrolase domain (HD), which decreases cellular dNTP levels (13–15). The HD domain has also been shown to be required for protein oligomerization and RNA binding, and has been suggested to have nuclease activity, although this is a matter of debate (16–18).

Our interest in SAMHD1 was stimulated by the other known restriction roles this protein has in viral life cycles; SAMHD1 has been predominantly characterized as a host restriction factor for HIV (18, 19). The inhibitory mechanism against HIV-1 was first linked to the dNTPase activity of SAMHD1, which lowers the intracellular dNTP concentration below the level required for viral reverse transcription. It has been suggested that SAMHD1, or associated proteins TREX1 and RNaseH1, restricts HIV-1 through an RNase activity (16). However, the relative contribution of these two activities in viral restriction is still a matter of debate, and some reports challenge the presence of nuclease activity (17–19). In cycling cells, SAMHD1 is phosphorylated by cyclin-dependent kinase 1 or 2 (CDK1, CDK2) at threonine 592 (19). De-phosphorylation at residue T592 regulates the resistance to HIV-1 infection in non-cycling cells, such as myeloid cells or resting T cells (19, 20). However, whether the phosphorylation of SAMHD1 at residue T592 influences the enzymatic function of the protein remains controversial; it has been suggested that this residue does not influence the dNTPase function (21), whereas other studies propose that dNTPase-competent SAMHD1 homo-tetramers are destabilized through phosphorylation at T592 (22). Therefore, the precise role for T592 phosphorylation in regulating SAMHD1 function remains to be fully elucidated.

SAMHD1 is an interferon-stimulated gene (ISG). SAMHD1 function is elevated by interferon, suggesting that it could be involved broadly in antiviral defense (23). In support of this idea, SAMHD1 restricts not only HIV, but also DNA viruses (24–26). There is also evidence of viruses countering SAMHD1 restrictive activity. The HIV-2 encoded accessory factor Vpx, which is also found in closely related simian immunodeficiency virus (SIV) strains, degrades SAMHD1 through a proteasome-dependent mechanism (27). Our recent study has also shown that conserved herpesvirus protein kinases antagonize SAMHD1 restriction through phosphorylation (http://dx.doi.org/10.2139/ssrn.3255560). Downregulation of SAMHD1 expression by HPV16 could also counter the anti-viral activity of this protein.

In this report we confirm that SAMHD1 RNA and protein levels are downregulated by HPV16 in oral keratinocytes and that the E6 and E7 oncoproteins contribute to this downregulation. Overexpression of SAMHD1 had no effect on the HPV16 viral life cycle in oral keratinocytes, but it was notable that the exogenous SAMHD1 was not expressed in the differentiated epithelium therefore could not have any effect on viral genome amplification which occurs in the differentiated epithelium. CRISPR/Cas9 editing of SAMHD1 resulted in hyper-proliferation of basal cells during organotypic raft cultures of NOKs containing HPV16, but not in parental NOKs. This was confirmed by a “thickening” of the epithelium by HPV16 in the absence of SAMHD1, by elevated BrdU labeling of basal cells and enhanced expression of the S phase marker protein Cyclin E. There was also increased viral replication in the differentiated layer of the epithelium in the absence of SAMHD1. These results were obtained using 3 independent guide RNAs for CRISPR/Cas9 editing, and also in tonsil keratinocytes immortalized by HPV16. In proliferating monolayer cells CRISPR/Cas9 targeting of SAMHD1 had no effect on either cell proliferation of viral genome copy number. Therefore, the results present an intricate interplay between the virus and SAMHD1 that contributes to a controlled HPV16 life cycle only in differentiating epithelium. The results also demonstrate the utility of our NOKs system as it has allowed us to detect hyper-proliferation of epithelial cells in the absence of SAMHD1 only in the presence of HPV16, demonstrating a functional interaction between HPV16 and SAMHD1 that regulates cellular growth. This would be technically limiting using primary keratinocytes as it would be difficult to generate primary cells with SAMHD1 knocked out that would retain any proliferative capacity for differentiation studies. We propose that our system offers a unique model for identifying host proteins that specifically interact with HPV16 to regulate host cell growth and viral replication in oral keratinocytes.

## Results

### SAMHD1 is downregulated in HPV16 containing oral keratinocytes

Previous work from this lab developed and characterized an HPV16 life cycle model in TERT immortalized oral keratinocytes (NOKs) (9). Initial comparison of two clonal cell lines containing HPV16 that support late stages of the viral life cycle (NOKs+HPV16A and NOKS+HPV16B) with the parental NOKs revealed a decrease in both SAMHD1 RNA and protein expression (Figures 1A and 1B, respectively). In the presence of HPV16, SAMHD1 RNA is expressed at a 50% lower level than the NOKs (Figure 1A), which validates our previous observations in RNAseq analysis (9). Protein levels are decreased correspondingly (Figure 1B). This was consistent in three individual repeats, which were quantified (Figure 1B).

**Figure 1:**
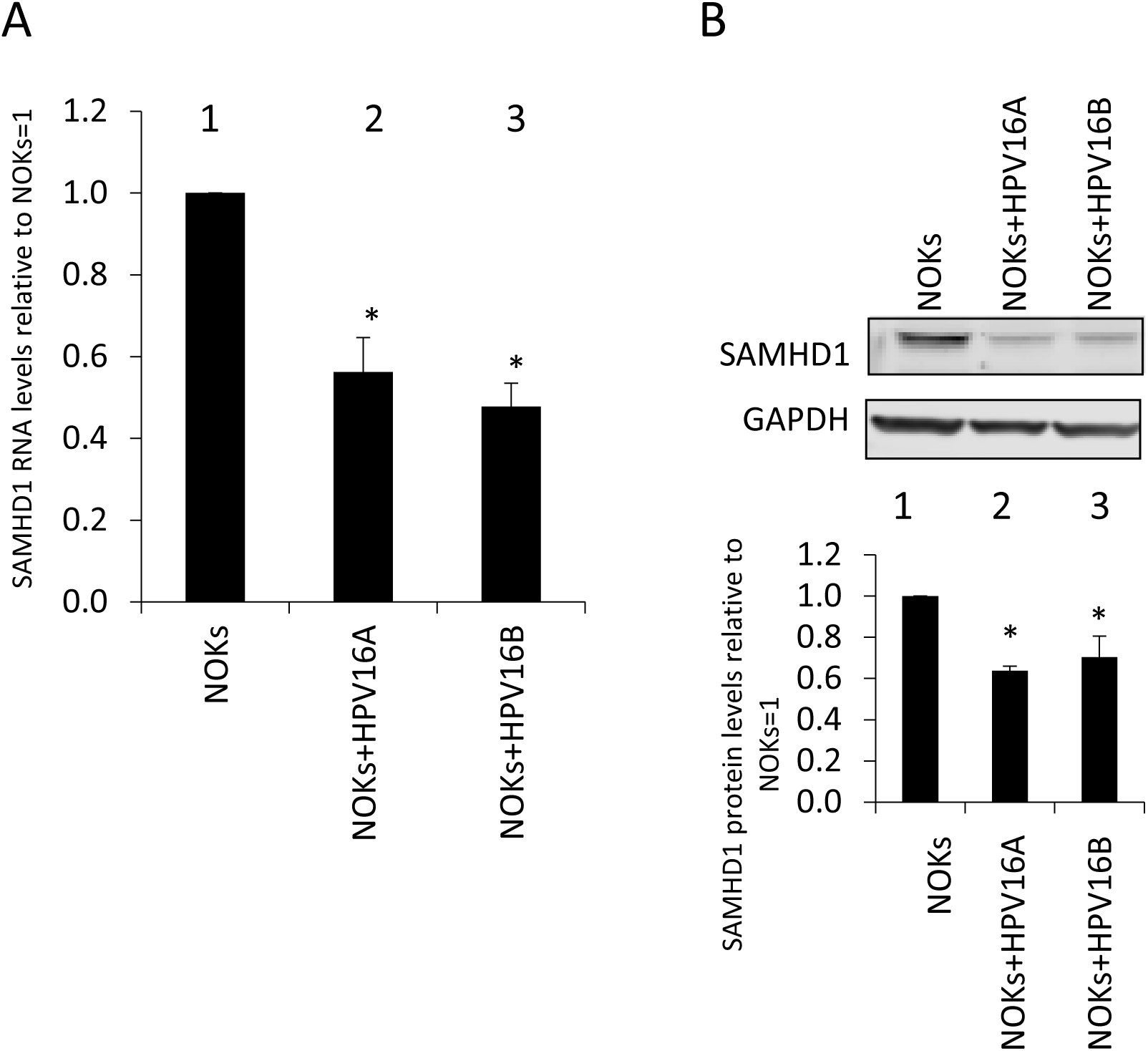
SAMHD1 is downregulated in HPV16 positive oral keratinocytes. A) RNA expression levels of SAMHD1 in NOKs (lane 1) and HPV16-containing NOKs (lanes 2 and 3). Results are expressed as fold change from that observed in parental NOKs and represent the average of three independent experiments. B) Western blot analysis was carried out on protein extracted from the NOKs (lane 1) and HPV16-containing NOKs (lanes 2 and 3). GAPDH is shown as an internal control. Western blots were visualized and quantitated using a Licor and calculated relative to parental NOKs. Data in A and B represents the average of 3 independent experiments and error bars indicate standard error of the mean. * indicates p<0.05.

To identify whether a single HPV16 protein is responsible for the downregulation of SAMHD1, stable cell lines were generated expressing individual viral proteins E6, E7 and expressing both oncoproteins together (NOKs+E6, NOKs+E7, NOKs+E6E7). Analysis of SAMHD1 expression in these cells reveals a downregulation at an RNA level in NOKs+E6E7, but not when the viral proteins are expressed individually (Figure 2A), suggesting a synergistic mechanism of action in SAMHD1 transcriptional repression. This decrease in RNA corresponded to a lower level of SAMHD1 protein in NOKs+E6E7 when compared with NOKs (Figure 2B). A downregulation in SAMHD1 protein was also observed in cells expressing single HPV16 oncoproteins, suggesting a post-transcriptional mechanism for downregulating SAMHD1 expression (Figure 2B). Therefore, E6 and E7 use a combination of transcriptional and post-transcriptional mechanisms to downregulate SAMHD1 expression.

**Figure 2:**
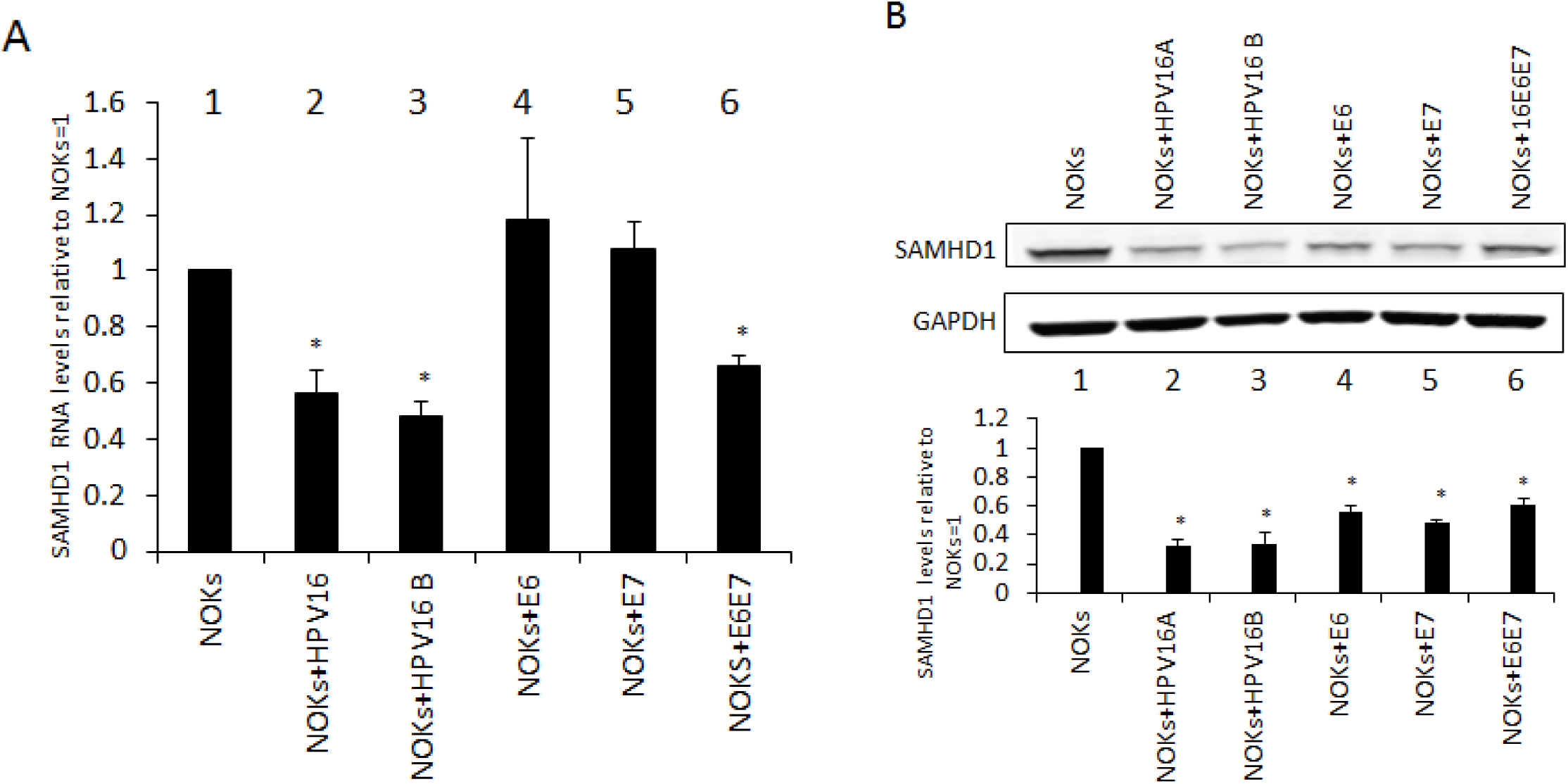
HPV16 oncoproteins affect SAMHD1 RNA and protein levels. A) RNA expression levels of SAMHD1 in NOKs (lane 1), HPV16-containing NOKs (lanes 2 and 3) and NOKs expressing HPV16 oncoproteins (lanes 4, 5 and 6). Results are expressed as fold change from that observed in parental NOKs and represent the average of three independent experiments. Western blot analysis was carried out on protein extracted from the NOKs (lane 1), HPV16-containing NOKs (lanes 2 and 3) and NOKs expressing HPV16 oncoproteins (lanes 4, 5 and 6). GAPDH is shown as an internal control. Western blots were visualized and quantitated using a Licor and calculated relative to parental NOKs. Data in A and B represents the average of 3 independent experiments and error bars indicate standard error of the mean. * indicates p<0.05.

### Downregulation of SAMHD1 by HPV16 is maintained during differentiation

In order to assess the expression of SAMHD1 during the HPV16 life cycle, NOKs and NOKs+HPV16 were differentiated by organotypic ‘raft’ culture. The rafts were then fixed and subject to immunofluorescent staining to determine SAMHD1 levels and localization in differentiated epithelia. SAMHD1 is expressed in NOKs throughout the organotypic section (Figure 3A), whereas fewer cells are stained when HPV16 is present (NOKs+HPV16A and NOKS+HPV16B). Three independent organotypic raft cultures were stained and SAMHD1 levels quantitated using a Vectra^®^ Polaris™ automated imaging system; the difference in staining between HPV negative and positive NOKs cells is significant (Figure 3B). Furthermore, the presence of HPV16 in NOKs leads to the loss of SAMHD1 expression in the upper layers of the epithelium (Figure 3C). This was quantified by measuring the “height” to which SAMHD1 is expressed in the rafts using a Vectra^®^ Polaris™ automated imaging system; in the case of HPV16 positive NOKs raft sections, SAMHD1 expression occurs from the basal layer to half way up the raft, while in NOKs staining is observed throughout the differentiated epithelium. It is also clear that the intensity of the SAMHD1 staining is diminished in the presence of HPV16 in NOKs (Figure 3A). Human tonsil keratinocytes immortalized by HPV16 (HTK16 in Figure 3) were also stained for SAMHD1 and again it is clear that, in the upper layers of the differentiated epithelium, there is no detectable levels of SAMHD1 staining (Figures 3A and 3C). These results confirm that the expression of SAMHD1 is downregulated by HPV16 during the viral life cycle in differentiating oral keratinocytes containing HPV16.

**Figure 3.**
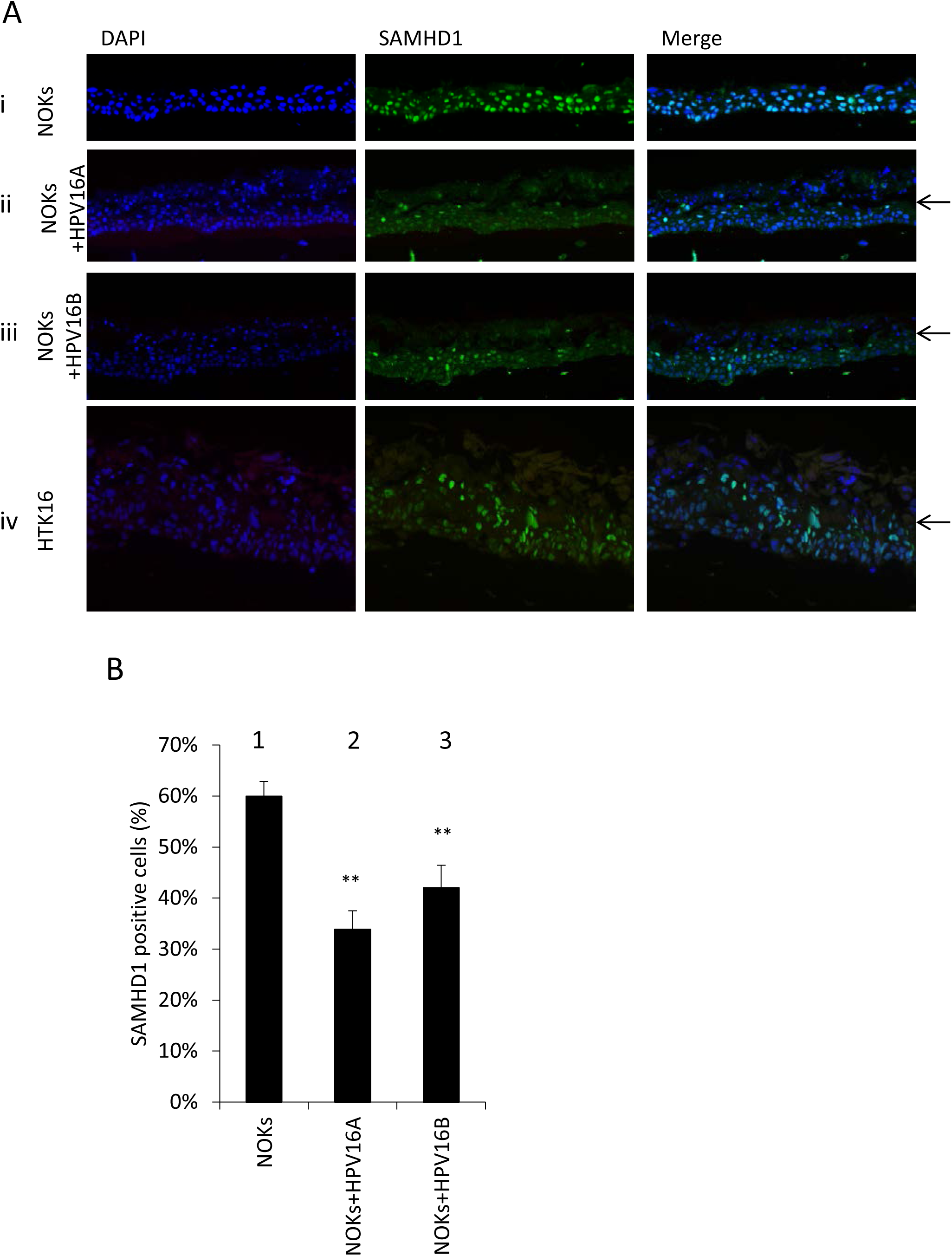
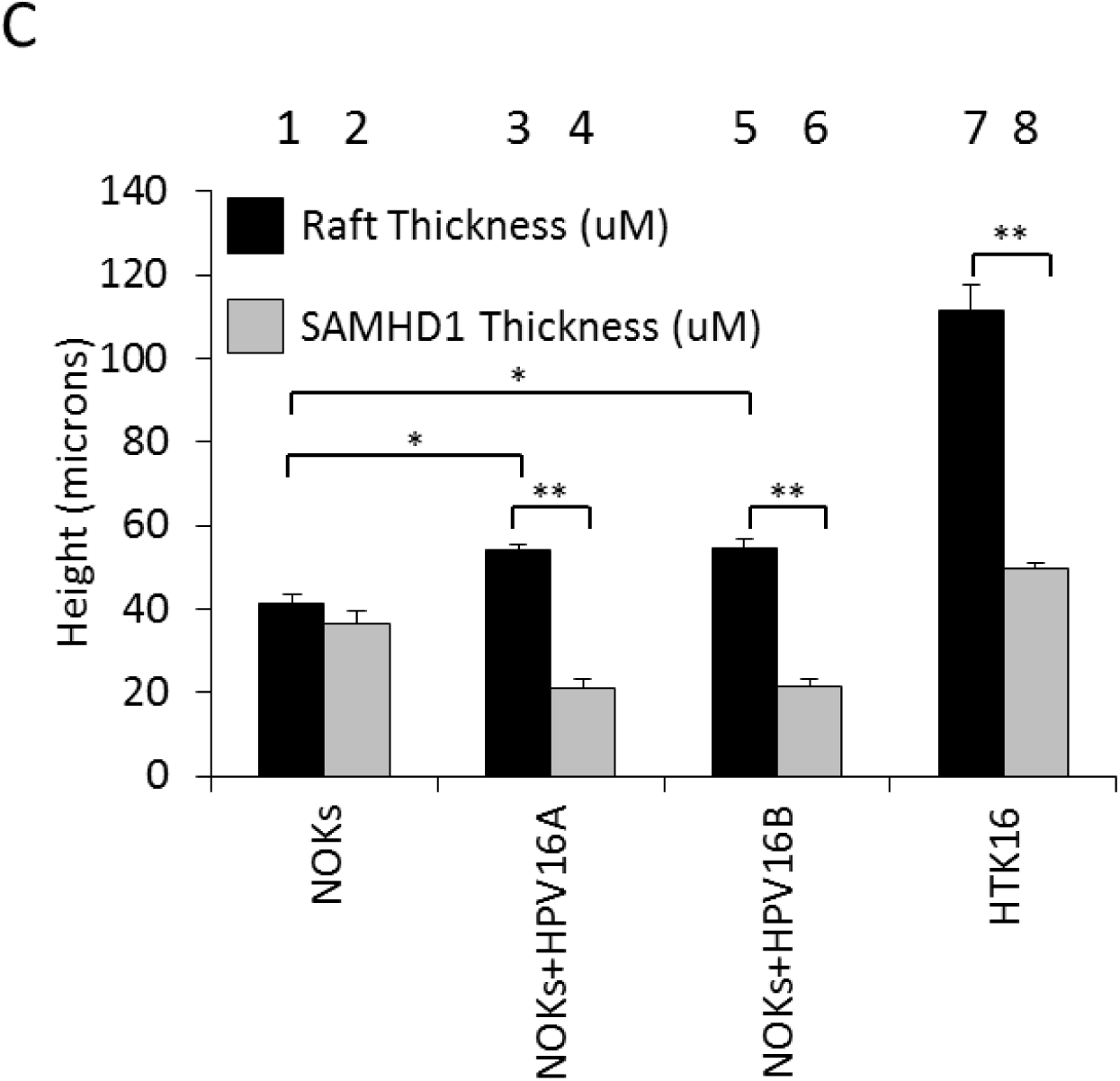
SAMHD1 is downregulated by HPV16 in differentiated epithelia. A) NOKs (i), NOKs+HPV16A (ii), NOKs+HPV16B (iii) and human tonsil keratinocytes immortalized by HPV16 (HTK16, iv), were differentiated in culture, and resulting sections stained for SAMHD1. Arrows indicate level at which SAMHD1 disappears. B) SAMHD1 positive cells of three sections from two individual rafts were quantified computationally and the measurements averaged. Immunofluorescence was quantified, using a Vectra^®^ Polaris™ automated imaging system, where whole stained sections were scanned computationally and the intensity calculated compared to a negative background control (secondary antibody only) and a positive localization control (DAPI). The same imaging parameters were used for each slide. For each sample, two sections from three individual rafts were scanned, to generate average values shown in this graph. Error bars indicate standard error of the mean and * indicates p<0.05. C)The overall thickness and region of SAMHD1 positivity from three sections from two individual rafts was quantified computationally using a Vectra^®^ Polaris™ automated imaging system and the measurements averaged. Error bars indicate standard error of the mean and * indicates p<0.05.

### Exogenously expressed SAMHD1 remains downregulated in the differentiated epithelium of HPV16 positive cells

The downregulation of SAMHD1 may promote the HPV16 life cycle as this would increase the nucleotide pool in the epithelium that could assist with viral genome replication. We therefore generated NOKs, NOKs+HPV16 (A and B clones) and HTK16 with exogenously overexpressed SAMHD1 to investigate the ability of this excess SAMHD1 to block genome amplification (Figure 4A). These cells were subjected to organotypic raft cultures and then fixed and stained with a V5 antibody (Figure 4B). The exogenous V5 tagged SAMHD1, like endogenous SAMHD1 (Figure 3), is not detectable in the upper layers of the differentiated epithelium when HPV16 is present. This was repeated and quantitated using a Vectra^®^ Polaris™ automated imaging system (Figure 4C). Due to the lack of SAMHD1 expression in the upper layers of the differentiated epithelium there was no change in the viral life cycle in the cells over expressing V5 tagged SAMHD1 (Figure S1).

**Figure 4:**
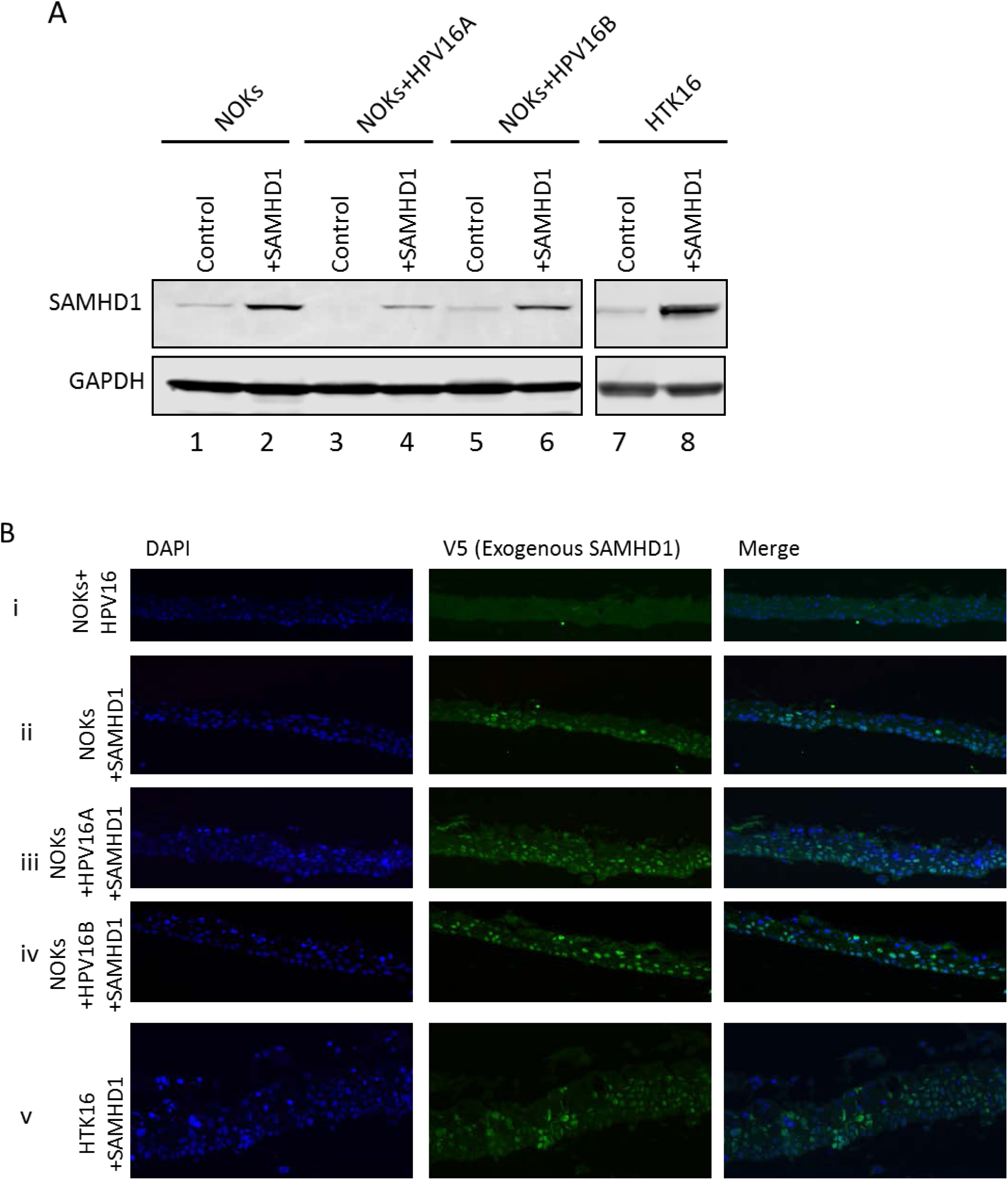
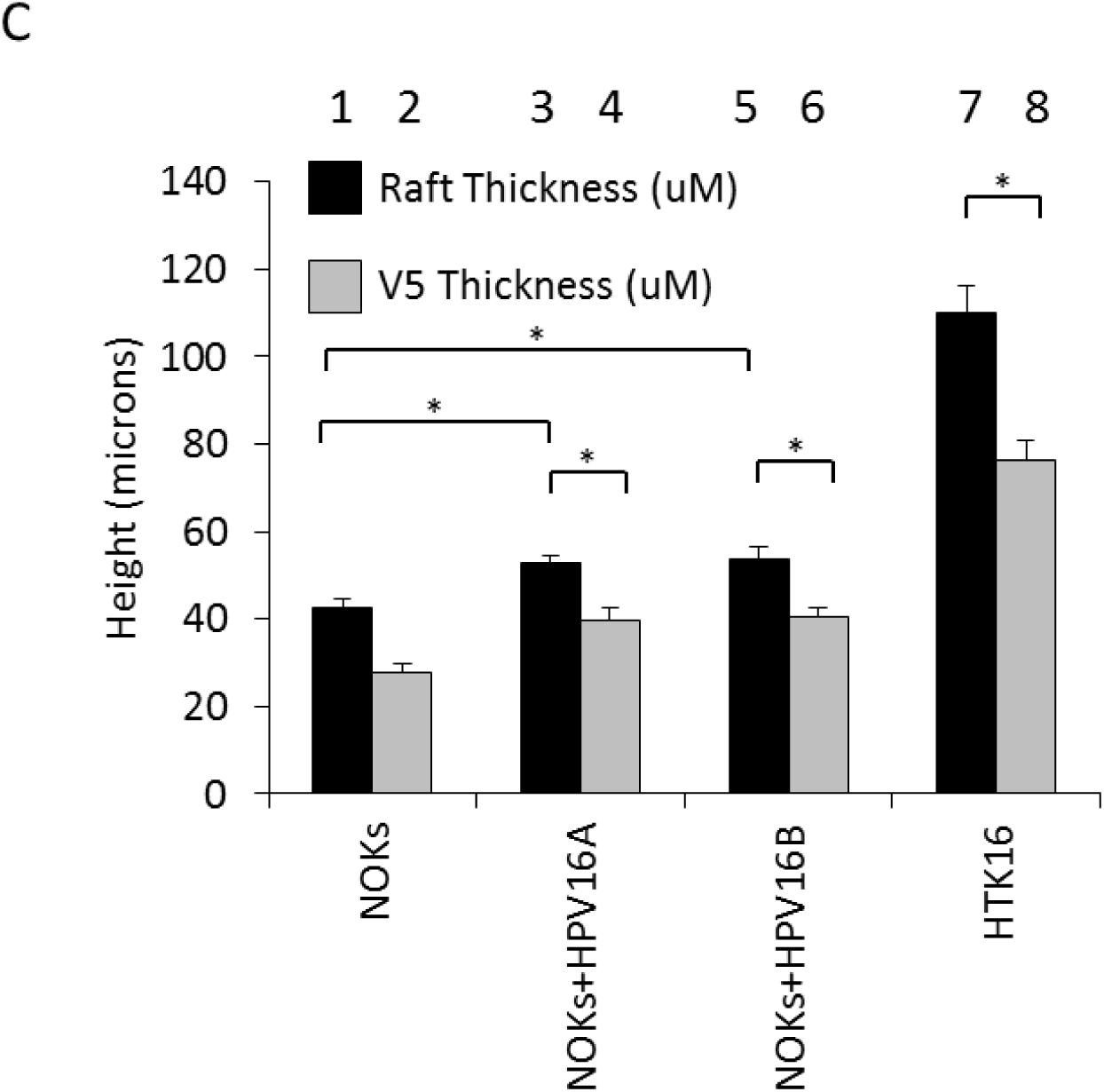
V5-tagged SAMHD1 is not expressed in the upper layers of epithelium during differentiation. A) NOKs, NOKs+HPV16A and NOKs+HPV16B and HTK16 cells overexpressing V5-tagged SAMHD1 were generated by transfection of cells with pLX304-SAMHD1 and selection with blasticidin. Expression of V5-tagged SAMHD1 in these cell lines was confirmed by western blot. HTK16 samples (lanes 7 and 8) were run on a separate gel. All of the control cells have been selected following infection with empty pLX304. B) Representative images of raft sections stained for exogenous SAMHD1, using an anti-V5 antibody. C) Sections were stained for exogenous SAMHD1, using an anti-V5 antibody. Immunofluorescence from three sections each from two individual differentiated cultures was then quantified using a Vectra^®^ Polaris™ automated imaging system and the measurements averaged. Error bars indicate standard error of the mean and * indicates p<0.05.

### Deletion of SAMHD1 results in hyper-proliferation of HPV16 positive keratinocytes in organotypic raft cultures

While HPV16 downregulates the expression levels of SAMHD1 in keratinocytes (Figures 1 and 2), there remains a substantial level of SAMHD1 in the HPV16 positive cells. To determine whether this remaining SAMHD1 played an important role in regulating the HPV16 life cycle in keratinocytes, SAMHD1 expression was removed using CRISPR/Cas9 targeting. Figure 5A demonstrates successful downregulation of SAMHD1 protein in the targeted cells. These were pools of cells therefore there remains a residual level of SAMHD1 expressed in cells not successfully targeted by CRISPR/Cas9. Having established stable knockdowns, these cells were differentiated via organotypic raft culture. Initial H&E staining revealed a hyper-proliferative phenotype in epithelia where SAMHD1 was knocked down and HPV16 was present, compared to control cells (NOKs by themselves) and to cells overexpressing SAMHD1 that were described in Figure 4 (Figure 5B). Triplicate rafts were sectioned and stained with H&E, before measurement using a Vectra^®^ Polaris™ automated imaging system. Quantification revealed that this increase in section thickness by HPV16 was significant (Figure 5C). This hyperproliferation was not due to the SAMHD1 depleted cell lines growing quicker in monolayer cells (Figure 5D). Additionally, there was no difference in HPV16 genome copy number in monolayer cells that under express SAMHD1 (Figures 5E&5F).

**Figure 5.**
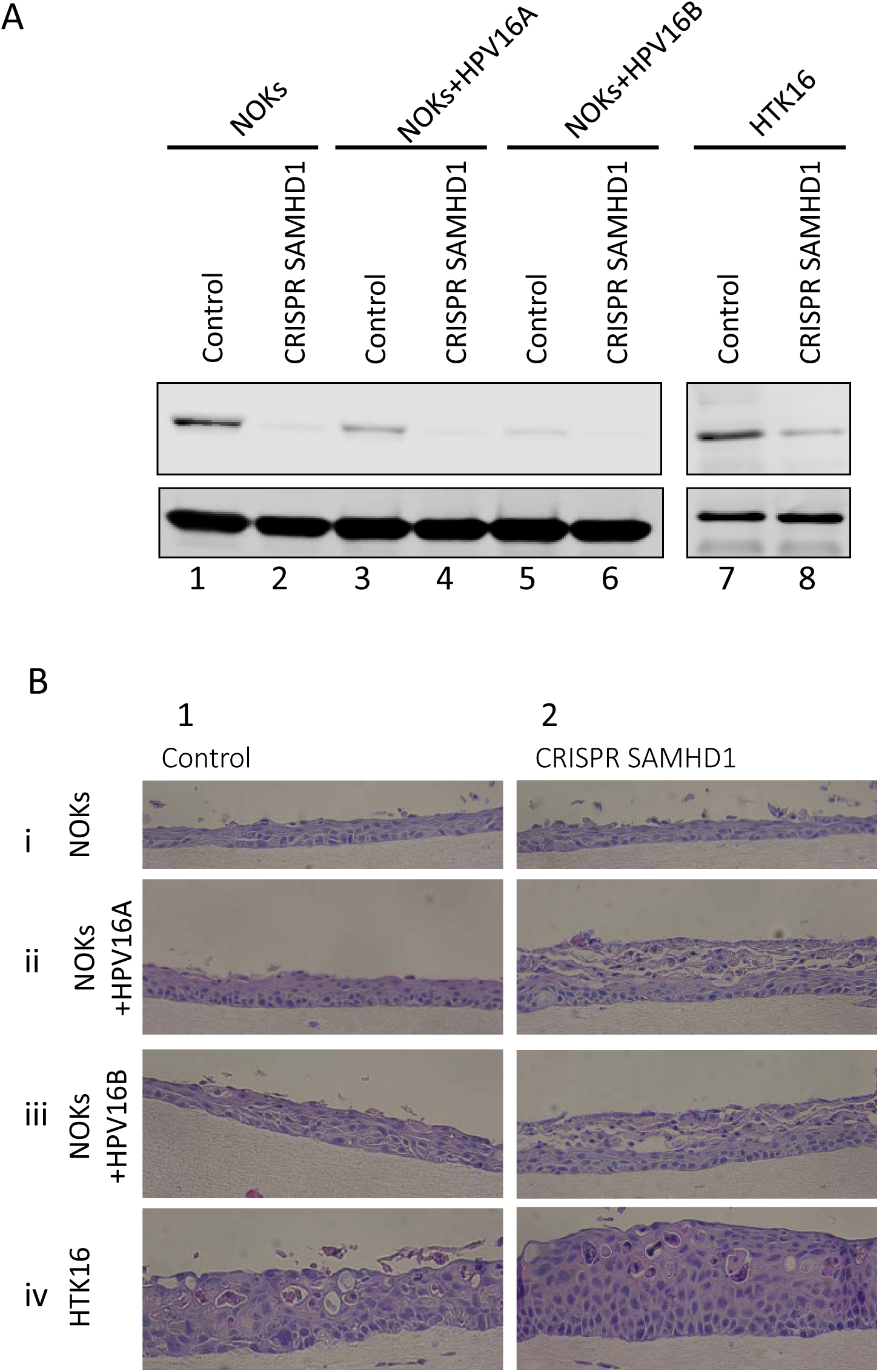
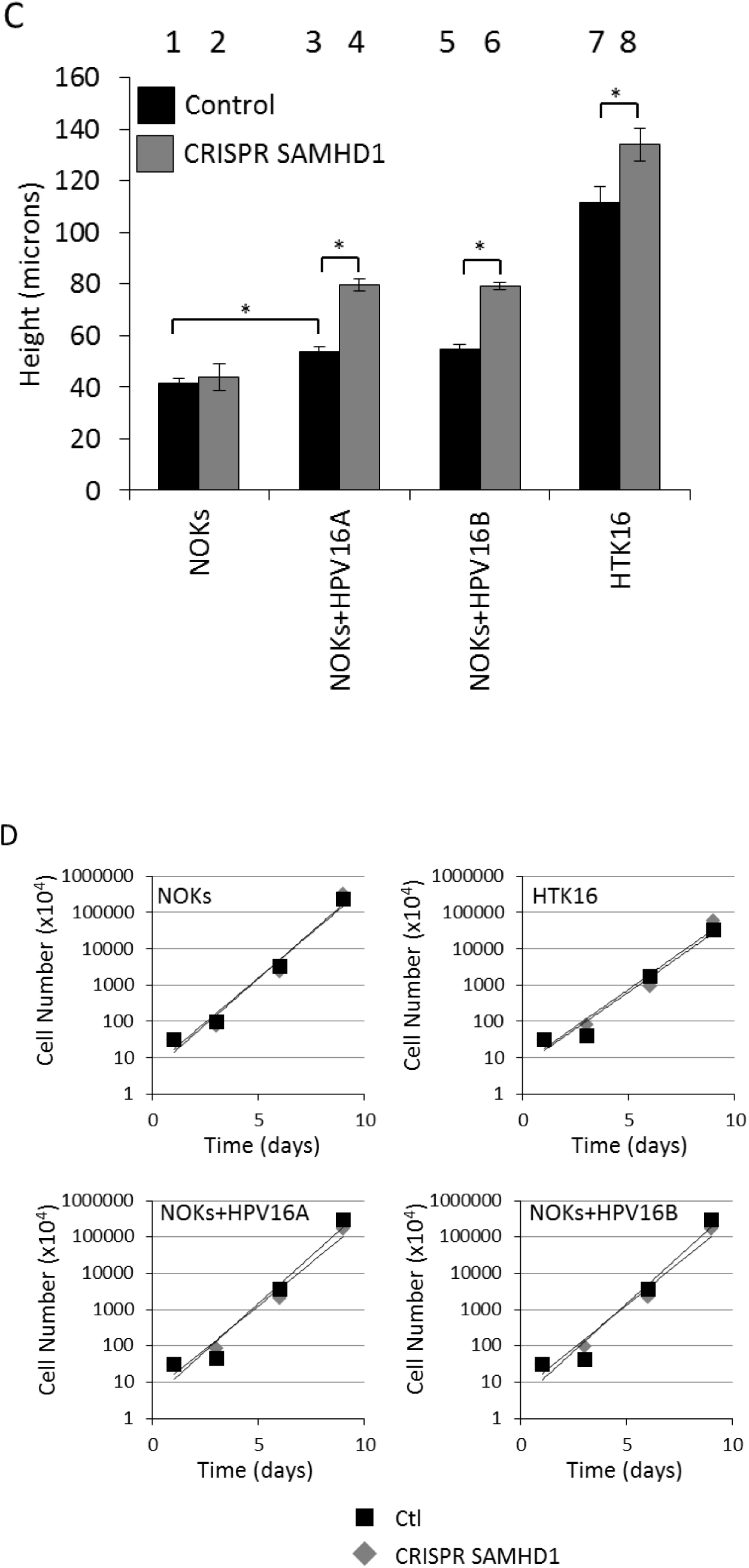
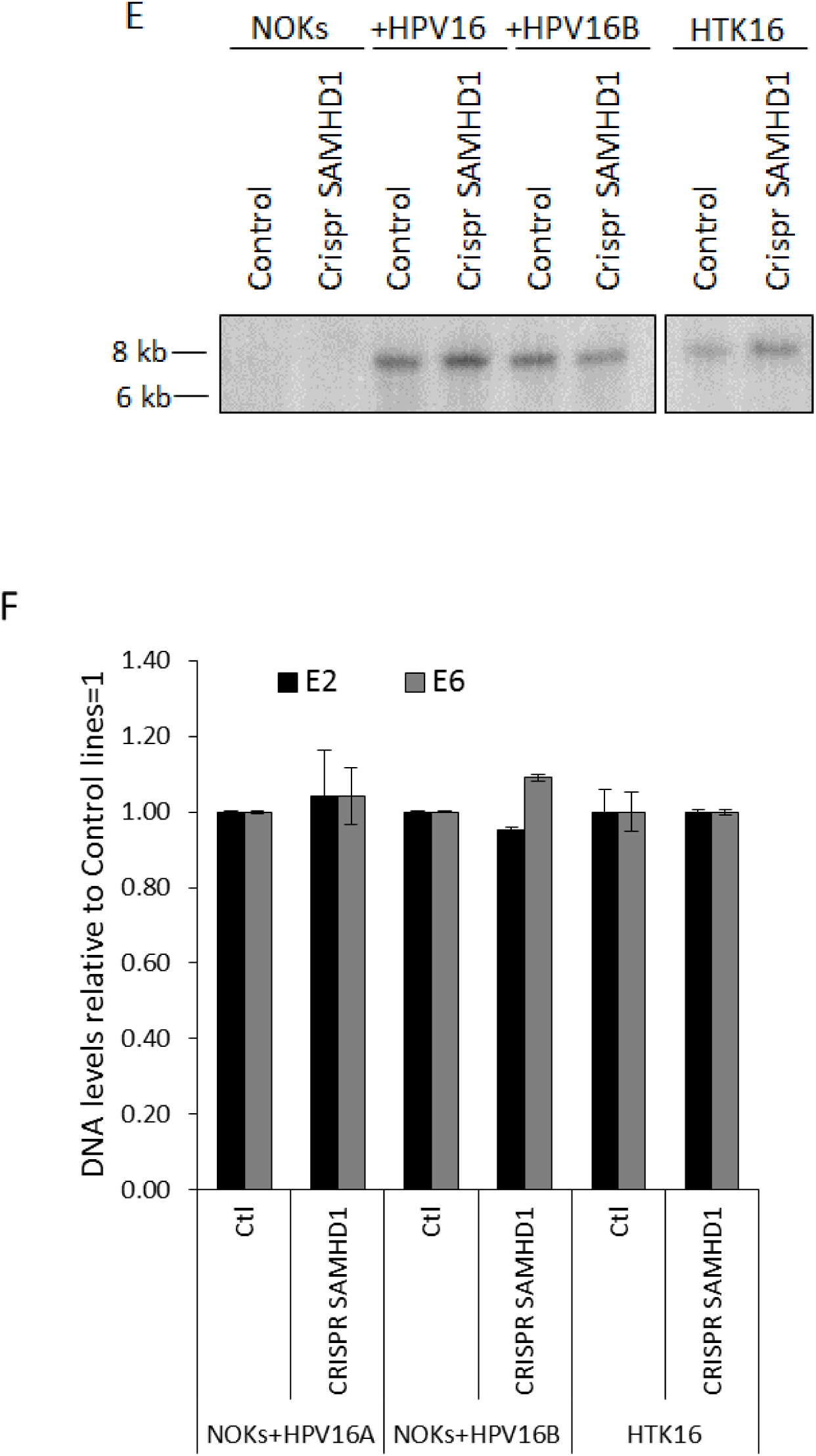
Generation and preliminary analysis of SAMHD1 CRISPR NOKs. SAMHD1 knockdown cell lines were generated in NOKs, NOKs+HPV16A and HTK16 cells by infection of cells with lentiviruses containing SAMHD1 CRISPR guide RNAs, followed by selection with puromycin. Depletion of SAMHD1 was confirmed by western blot; samples from HTK16 (lanes 7 and 8) were run on a separate gel. Control cells were generated identically but with control guide RNA. B) Representative images of rafts that were quantified. When compared to compared to parental lines (column 1), SAMHD1 CRISPR lines (column2) appeared to be hyper proliferative where HPV16 was present. C) Three H and E stained sections from two individual rafts were imaged and measurements were taken at 100 micron intervals across each section using a Vectra^®^ Polaris™ automated imaging system and quantitated. Error bars indicate standard error of the mean and * indicates p<0.05. D) SAMHD1 CRISPR cell lines were plated alongside parental cell lines at 3×10^5^, grown for 3 days before harvesting and counting. This was repeated three times to assess cell growth rates in monolayer grown cells, which are unaltered by the depletion of SAMHD1. E) Southern blots were carried out on DNA extracted from the indicated cell lines. The DNA was digested with Sph1 to linearize the 8kbp viral genome and the resultant blot was probed with the labeled HPV16 DNA. F) DNA was extracted from control and SAMHD1 CRISPR cells lines and subject to PCR detection of HPV16 E2 and HPV16 E6. DNA from NOKs was utilized as a negative control. Triplicate samples were analyzed and averaged, error bars indicate standard error of the mean.

To investigate whether this hyper-proliferation was specific to a certain layer of the differentiated culture, rafts were treated with BrdU for the final 16 hours of culture, before being fixed and sectioned. BrdU staining highlights the cells actively dividing in those 16 hours. In NOKs cells there was no increase in BrdU positive cells in the absence of SAMHD1, Figure 6A. In NOKs+HPV16 (A and B clones) there was an increase in BrdU labeling compared with NOKS cells, demonstrating the expected enhanced proliferation induced by HPV16 (Figure 6A). In addition, when SAMHD1 was removed there was a further increase in BrdU labeling in the presence of HPV16 that is not observed in the parental NOKs. There was no disruption in the differentiation of these cells in the absence of SAMHD1 as determined by involucrin staining (Figure 6A). These experiments were repeated with HTK16 cells lacking SAMHD1 (Figure 6A v and vi) where, as with NOKs+HPV16, there is an increase in BrdU positive cells in the absence of SAMHD1. These experiments were repeated and quantitated using a Vectra^®^ Polaris™ automated imaging system (Figure 6B) and there is a statistically significant increased level of BrdU positive cells in NOKs+HPV16 and HTK16 but not in NOKs. This demonstrates an interaction between HPV16 and SAMHD1 that controls cellular proliferation in the basal layers of the epithelium.

**Figure 6.**
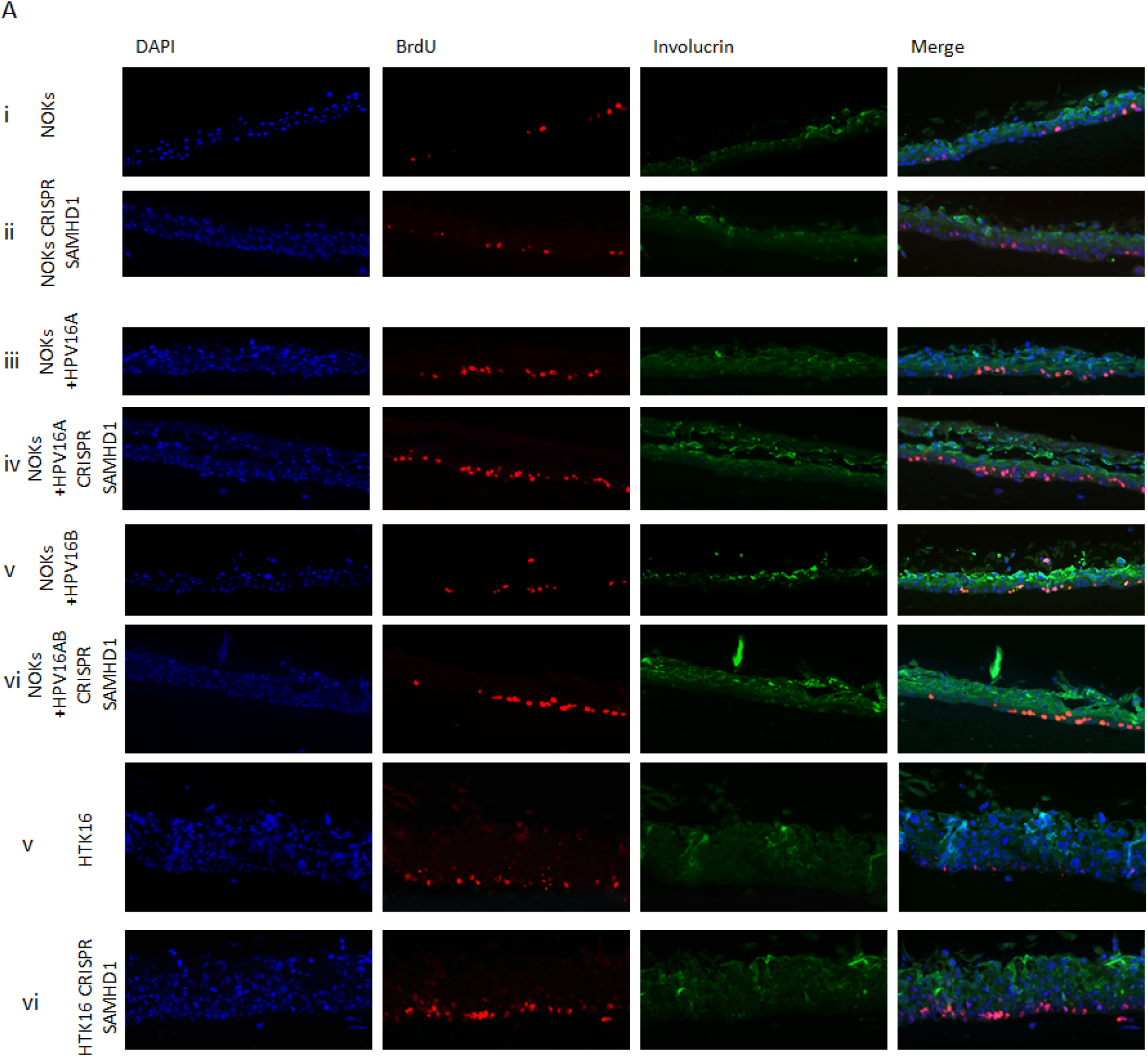
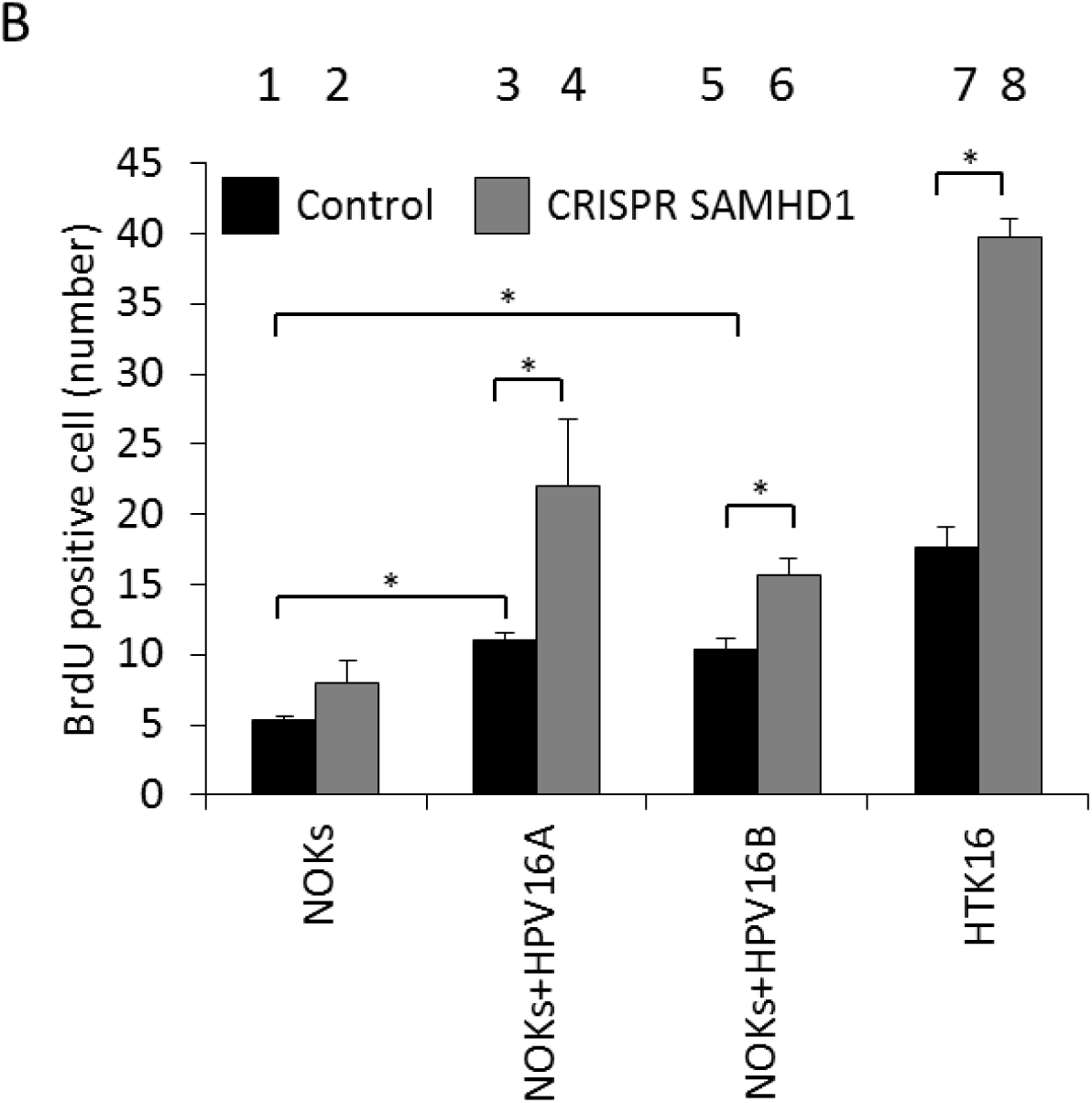
Knockdown of SAMHD1 increases proliferation in basal cells of HPV16 positive oral epithelia. A) Nucleoside analog BrdU was included in media for the final 16 hours of organotypic raft culture. Fixed sections were then stained for BrdU incorporation and for differentiation marker involucrin. B) The number of BrdU positive cells was measured using a Vectra^®^ Polaris™ Imaging System, whereby whole stained sections were scanned computationally and the intensity calculated compared to a negative background control (secondary antibody only) and a positive localisation control (DAPI). The same imaging parameters were used for each slide. Three sections from two individual rafts were subject to analysis, error bars indicate standard error of the mean and * indicates p<0.05.

To further confirm the proliferative nature of these cells, cells were stained with Cyclin E, a protein expressed in S phase (28). In NOKs cells there is no increase in Cyclin E staining without SAMHD1, while there is increased Cyclin E staining in the absence of SAMHD1 in NOKs+HPV16A (Figure 7A). This increase in Cyclin E staining in the absence of SAMHD1 was also observed in HTK16 cells lacking SAMHD1 when compared with parental HTK16. These results were repeated and Cyclin E positive cells quantitated using a Vectra^®^ Polaris™ automated imaging system and a summary of the results is presented graphically in Figure 7B. In NOKs+HPV16 and HTK16 there is a statistically significant increase in Cyclin E expression in the absence of SAMHD1 while no such increase is observed in NOKs.

**Figure 7.**
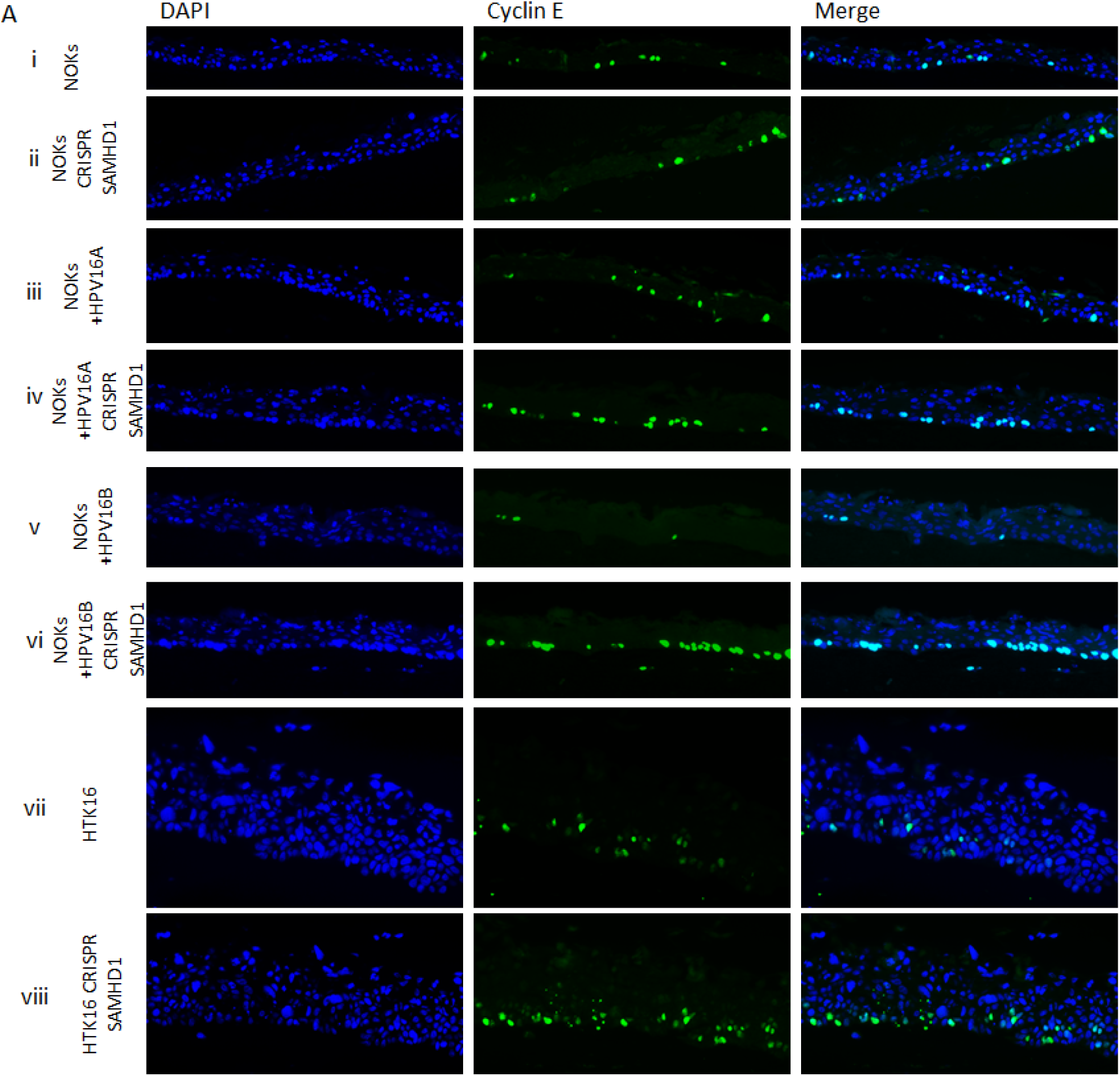
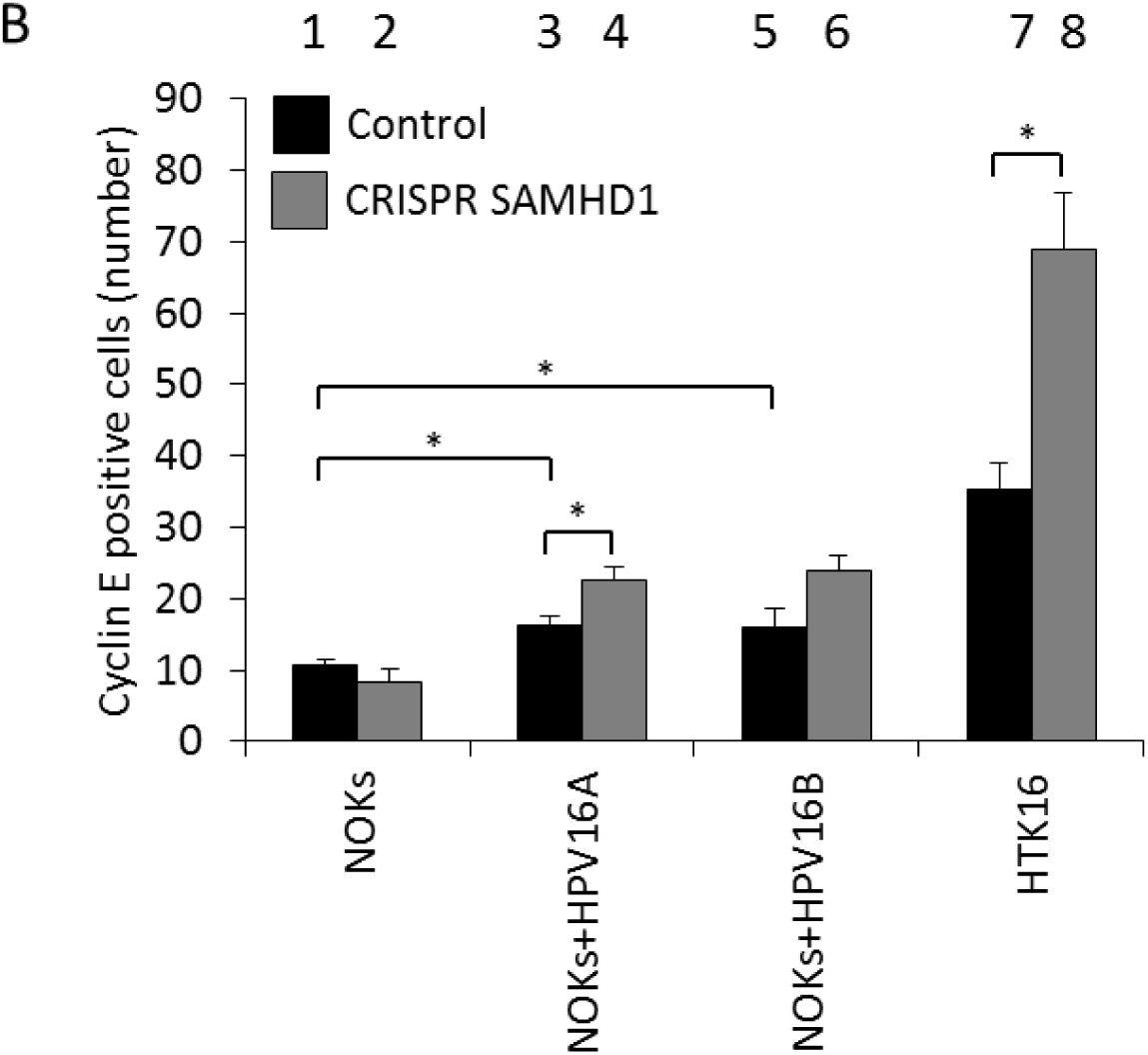
Knockdown of SAMHD1 increases proliferation in basal cells of HPV16 positive oral epithelia. A) Differentiated sections from organotypic raft cultures were stained for S phase marker Cyclin E. Representative images of Cyclin E staining in differentiated NOKs, NOKs+HPV16 and HTK16 cell lines is shown. B) The number of Cyclin E positive cells was counted was measured Vectra^®^ Polaris™ Imaging System, whereby whole stained sections were scanned computationally and the intensity calculated compared to a negative background control (secondary antibody only) and a positive localization control (DAPI), using the same imaging parameters for each slide. Three sections from two individual rafts were subject to analysis, error bars indicate standard error of the mean and * indicates p<0.05.

As there was a hyper-proliferation of the HPV16 positive cells in the absence of SAMHD1, and an increased thickening of the differentiated epithelium, we investigated whether there was an enhanced amplification of the HPV16 genome in the cells lacking SAMHD1. To do this FISH analysis for the viral DNA was carried out and the intensity of the staining was measured using the Vectra^®^ Polaris™ automated imaging system; whole stained sections were scanned computationally and the intensity and localization of staining measured relative to a negative control (NOKs) and a positive control (NOKs+HPV16). Figure 8A shows representative images from the staining. Figure 8B summarizes the results of these experiments and in both NOKs clones that have HPV16 and SAMHD1 removed there is a statistically significant enhanced FISH signal detected indicating increased HPV16 replication. There was also an increased signal in HTK16 in the absence of SAMHD1. In 8Ai it is clear that there is no signal in the NOKs while in 8Aii and 8Aiii there is an enhanced FISH signal in the absence of SAMHD1. In HTK16 (8Aiv) there is also a slight increase in signal. Please note that the measurement of the signal is quantitative and non-subjective while the images are representative. There was a hint of viral genome amplification in the basal layers of the HPV16 containing cells in the absence of SAMHD1. In Figure 8C we have increased the exposure and it is clear that there is some HPV16 DNA signal in the absence of SAMHD1 prior to the amplification stage of the viral life cycle. This was difficult to quantitate but does suggest that viral genome replication is higher even in the basal layers of the differentiating epithelium.

**Figure 8.**
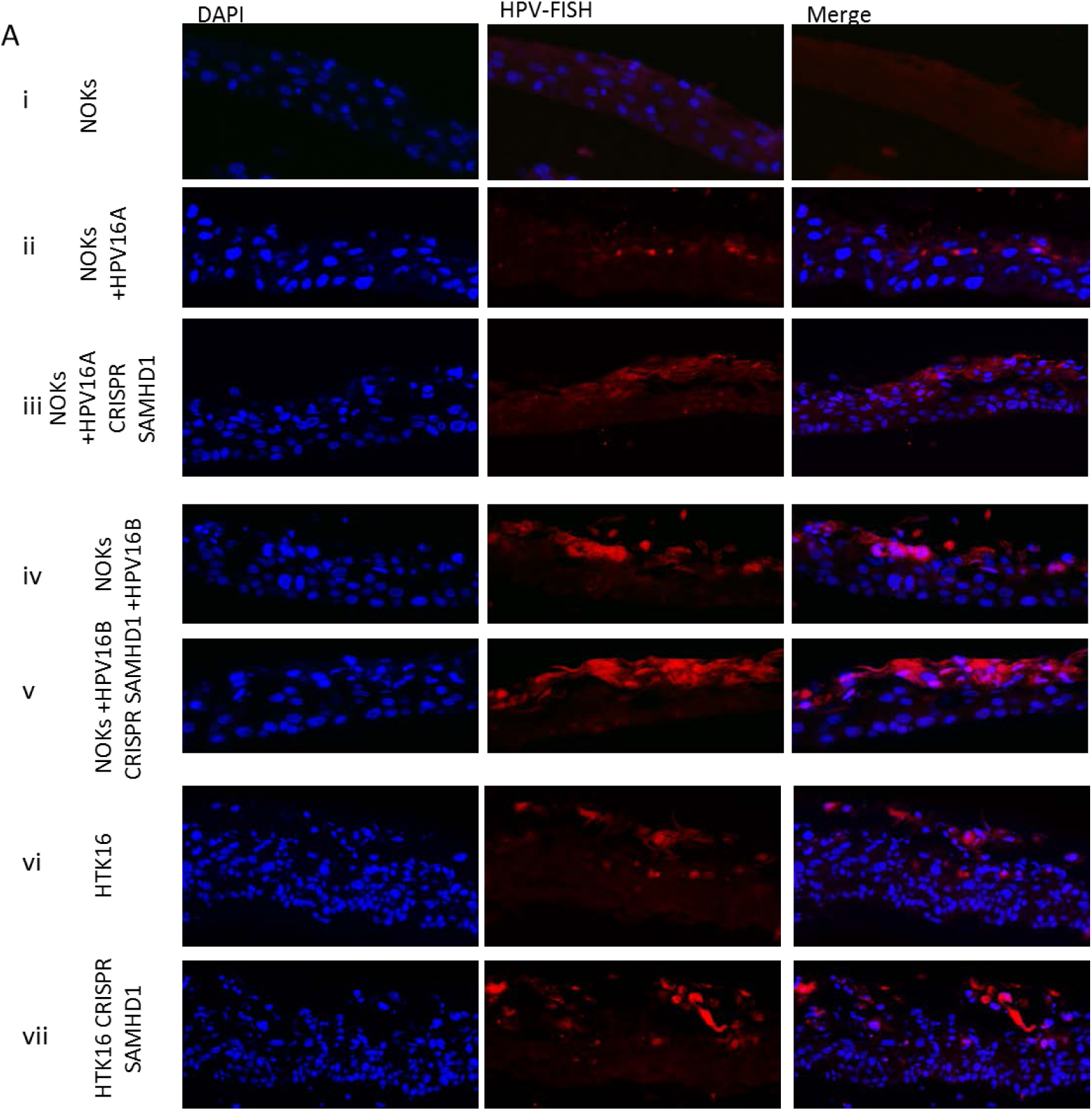
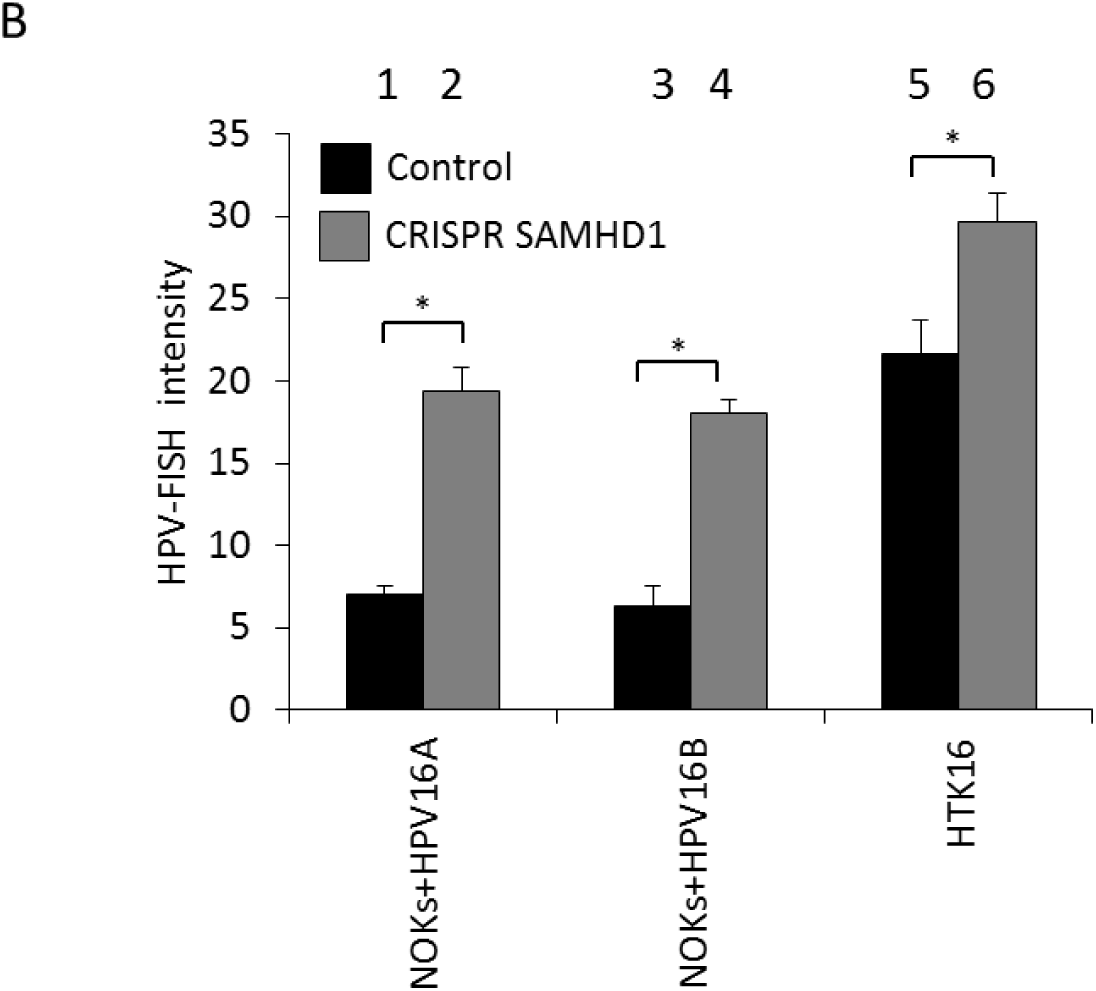
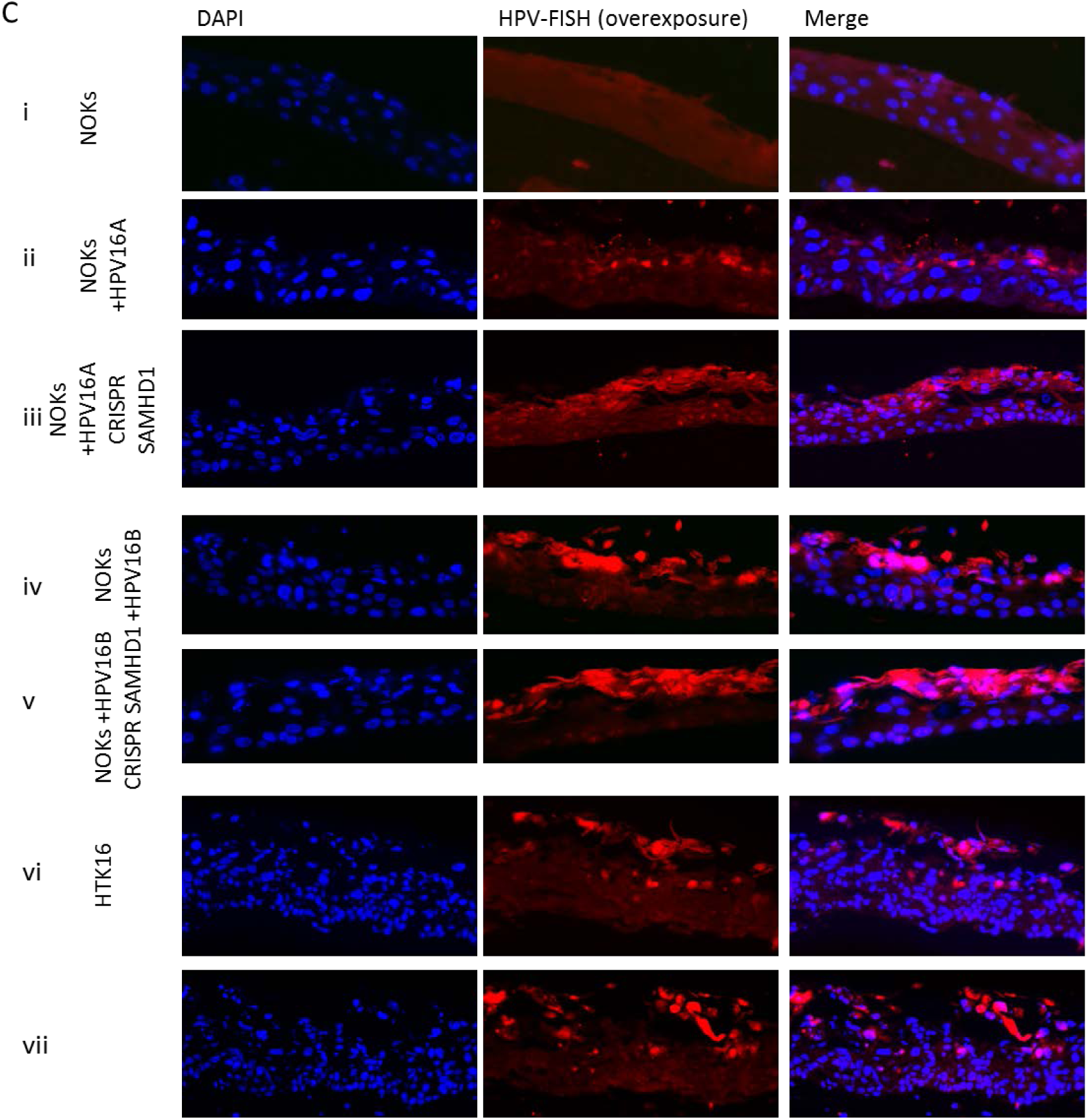
Knockdown of SAMHD1 increases HPV16 amplification in differentiated culture. A) NOKs+HPV16A and HTK16s were differentiated in culture, formalin fixed, paraffin embedded, sectioned and then stained HPV16 genomes using DNA-FISH. Images shown are representative images of HPV-FISH in differentiated culture. B) The intensity of fluorescence was quantified using Vectra^®^ Polaris™ Imaging System, whereby whole sections were scanned computationally and the intensity calculated compared to a negative background control (NOKs). C) An over exposure of the image shown in A to indicate viral genome detection in the basal layers in the absence of SAMHD1. The same imaging parameters were used for each slide in A and C. Error bars indicate standard error of the mean and * indicates p<0.05.

Finally, we reproduced our results with an additional two SAMHD1 guide RNAs that target different sequences in the SAMHD1 gene. Figure S1A details the knock down of SAMHD1 expression by these additional CRISPR/Cas9 targeting sequences. Figure S1B demonstrates a thickening of the epithelium in NOKs only in the presence of HPV16 following SAMHD1 removal while Figures S1C and S1D demonstrate the enhanced BrdU and Cyclin E staining, respectively, when SAMHD1 is removed in the presence of HPV16. Overall the results with the three independent SAMHD1 guide RNAs are identical demonstrating that the results are not the due to off target effects of the SAMHD1 guide RNAs.

## Discussion

Many excellent studies have been carried out with primary human keratinocytes to characterize the immortalization properties of high risk HPV (HR-HPV) These HPV immortalized cells have proved invaluable at identifying cellular proteins that are required for the life cycle of HR-HPV (3–6, 29–43). For example, it is clear that host cell homologous recombination factors are required for the amplification phase of the viral life cycle (5, 39, 40). Immortalized primary epithelial cells remain a crucial resource for characterization of host factors that regulate HR-HPV life cycles as well as investigating the immortalization properties of viral mutants. However, there are inevitably some limitations to using HPV immortalized keratinocytes. For example, if genes are knocked down to investigate the role in the viral life cycle, it is difficult to investigate what the consequences of this knock down are in non-HPV immortalized cells. This is because selection of primary cells with host genes knocked down takes several cell passages and primary cells will often senesce prior to selection. Even if knock-down cells were generated, they would have a very limited life span for making organotypic rafts to investigate the consequences of the gene knock down on normal cell differentiation. Given the current epidemic of HPV16+OPC we wanted to develop an HPV16 oral keratinocyte life cycle model that would have additional utilities to HPV16 immortalized primary cells. To do this we used NOKs which are immortalized by telomerase. We generated clonal cell lines of NOKs that retain episomal HPV16 genomes and demonstrated transcriptional reprogramming by HPV16 in these cells that is related to that regulated in HPV16 positive head and neck cancers. These NOKs+HPV16 cells, upon organotypic raft culture, demonstrate several markers of the late stages of the viral life cycle including E1^E4 expression, E2 expression and amplification of the viral genome in the upper layers of the differentiated epithelium (9).

Recently we demonstrated one potential unique use of this system. We expressed the HPV16 E2 protein in NOKs and determined host gene regulation by E2 in these cells that had a highly significant overlap with the genes regulated by the entire HPV16 genome (44). Therefore, this system has allowed us to identify and characterize a new function for the E2 protein; regulation of host gene transcription during infection. This study would not have been possible in primary keratinocytes as E2 would not immortalize these cells during selection. Importantly, in our studies using this system, we have validated gene expression changes in human tonsil keratinocytes immortalized by HPV16 (HTK16) and also in HPV16 positive head and neck cancers.

Here we report another utility of this system. We demonstrate that removal of SAMHD1 from NOKs+HPV16 and HTK16 results in hyper-proliferation of these cells in organotypic raft cultures. This increase in proliferation was confirmed by a “thickening” of the raft culture, and an increase in BrdU and Cyclin E positive cells in the absence of SAMHD1 (both S phase markers). We also observed an elevation in viral genome amplification in the absence of SAMHD1. The parental NOKs cells exhibit no increase in BrdU or Cyclin E positive cells, demonstrating that the increased proliferation was due to the presence of HPV16. If we had carried out these studies in the absence of the NOKs cells, then we would not have been able to determine whether the deletion of SAMHD1 expression by itself was proliferative for the cells. Therefore, this system has allowed us to identify an interaction between SAMHD1 and HPV16 that regulates the proliferation of NOKs+HPV16 and HTK16. The addition of HTK16 demonstrates that the SAMHD1 deletion phenotype is retained in cell lines immortalized by HPV16. Therefore, the combination of NOKs, NOKs+HPV16 and HTK16 represents an excellent system for identifying interactions between host proteins and HPV16 that have an effect on host cell growth and viral replication.

SAMHD1 is downregulated in NOKs+HPV16 and HTK16 and this is at least partially due to the viral oncogenes E6 and E7. As SAMHD1 is a dNTPase this downregulation could be important for promoting viral replication, particularly at the amplification stage of the viral life cycle in the differentiated epithelium. To counteract this down-regulation we overexpressed exogenous SAMHD1 but notably this protein was not expressed in the differentiated layers of the epithelium where genome amplification occurs. As E6 and E7 can both regulate SAMHD1 post-transcriptionally it is possible that the viral oncogenes remove SAMHD1 expression to allow viral genome amplification.

SAMHD1 is also a homologous recombination (HR) factor and is involved in recruiting MRE11 to damaged DNA (45, 46). This role of SAMHD1 could also play an important role in its interaction with HPV16 as this virus recruits a host of HR factors to its replicating DNA and it is proposed that HPV16 using HR during viral replication in order to amplify its genome. However, unlike downregulation of other HR factors (5, 39, 40), downregulation of SAMHD1 boosts viral genome amplification and also disrupts the equilibrium between the virus and the host cell.

The reason for the hyper-proliferation of HPV16 containing keratinocytes in the absence of SAMHD1 is not clear. This is not observed in monolayer cultures therefore only becomes apparent during organotypic raft cultures indicating that there may be some involvement of the collagen-fibroblast plug used to generate the differentiating cells. This collagen-fibroblast plug could mimic a stroma-epithelial cell interaction and there is a known cross-talk between HPV and the stroma (47). It is also apparent that there is not an increase in HPV16 genomes in the absence of SAMHD1 in monolayer cells so the difference is not due to an initial increased viral genome copy number in these cells upon rafting.

SAMHD1 is a restriction factor for HIV and other DNA viruses including HBV (26), HSV1 (24) and EBV (http://dx.doi.org/10.2139/ssrn.3255560) and here we demonstrate that SAMHD1 is also a restriction factor for HPV16. Not only does the absence of SAMHD1 promote hyper-proliferation of the infected cells, it also stimulates an enhanced amplification of the HPV16 genome during the terminal stages of differentiation. In addition, the FISH staining for the HPV16 genome in Figure 8C suggests that in NOKs+HPV16 in the absence of SAMHD1 there is an increase in viral signal in the basal layers of the epithelium. There is also an indication of this in HTK16. This is hard to quantitate but does suggest that during differentiation the increased replication of the viral genome perhaps starts early in the differentiation process.

Our isogenic NOKs system for investigating the HPV16 life cycle has been essential at revealing a specific interaction between the virus and SAMHD1 that controls host cell proliferation. In addition, the enhanced genome amplification in the absence of SAMHD1 demonstrates that SAMHD1 is a restriction factor for HPV16. The virus clearly downregulates SAMHD1 expression but retains a level that is required for controlling both host proliferation and viral genome amplification. Perhaps downregulation of SAMHD1 is required to generate an enhanced pool of nucleotides that would promote viral genome amplification, but that SAMHD1 homologous recombination function is also required to control the levels of viral genome replication. Future studies will focus on determining what structural and enzymatic functions of SAMHD1 contribute towards the control of HPV16 induced cellular proliferation, and what viral proteins SAMHD1 interacts with to regulate this control. Ultimately it may be possible to engineer elevated levels of functional SAMHD1 in the presence of HPV16 that could block HPV16 induced cellular proliferation and amplification of the viral genome.

## Materials and Methods

### Cell culture

Clonal cell lines containing the HPV16 genome were generated from normal oral keratinocytes (NOKs) as previously described (9). These cells were cultured alongside parental NOKs for all comparisons. NOKs and NOKs+HPV16 cells were grown in K-SFM (Invitrogen) with 1% (v/v) penicillin/streptomycin mixture (Thermo Fisher Scientific) containing 4 μg/mL hygromycin B (Millipore Sigma) at 37°C in a 5% CO_2_/95% air atmosphere and passaged every 3-4 days. NOKs+HPV16 were grown in the same medium also containing 150 μg/mL G418 (Thermo Fisher Scientific). NOKs expressing individual HPV16 oncoproteins were generated using a lentiviral delivery system, utilising the following expression plasmids: pMSCV16E6 (Addgene plasmid # 42603), pMSCV16E7 (Addgene plasmid # 35018) and pLXSN16E6E7 (Addgene plasmid # 52394). Resulting cell lines were cultivated in K-SFM with 1% (v/v) penicillin/streptomycin mixture, containing 2 μg/mL puromycin (HPV16 E6 and HPV16 E7) or 150 μg/mL G418 (HPV16 E6E7), at 37°C in a 5% CO2/95% air atmosphere and passaged every 3-4 days. Overexpression of SAMHD1 in NOKs, NOKs+HPV16 and HTK16 was achieved by lentiviral delivery of pLX304-SAMHD1 into cells and ensuing selection in 5μg/mL blasticidin (Millipore, Sigma). Similarly, CRISPR guides to SAMHD1 were delivered into NOKs, NOKs+HPV16 and HTK16 via lentivirus, and cells were selected by growth in puromycin containing k-SFM (2 μg/mL, Millipore Sigma). All cells were routinely checked for mycoplasma contamination. For downstream protein and RNA analysis, 1×10^6^ cells were plated onto 100 mm plates, trypsinized and pelleted after 24hrs and washed twice with phosphate buffered saline (PBS).

### SAMHD1depletion by CRISPR/Cas9 genome editing

CRISPR/Cas9-mediated SAMHD1 depletion was described previously (http://dx.doi.org/10.2139/ssrn.3255560). Briefly three different sgRNAs targeting human SAMHD1 were designed and cloned into lentiCRISPR v2 vector (Addgene plasmid # 52961). Packaging 293T cells were transfected with SAMHD1 sgRNAs (CRISPR SAMHD1) or a non-targeting sgRNA control (Control) and helper vectors (pMD2.G and psPAX2; Addgene plasmid #s 12259 and 12260 respectively) using Lipofectamine 2000 reagent (Cat# 11668019, Life Technologies). Medium containing lentiviral particles and 8 mg/mL polybrene (Sigma-Aldrich, St. Louis) was used to infect NOKs or HTK16 cells. Infected cells were selected in medium containing 2 μg/mL puromycin. The target guides sequences are as follows:

SAMHD1-sg1; F: 5’-caccgcttagttatatccagcgat-3’; R: 5’-aaacatcgctggatataactaagc-3’;

SAMHD1-sg2; F: 5’-caccgaatccacgttgatacaatga-3’; R: 5’-aaactcattgtatcaacgtggattc-3’;

SAMHD1-sg3; F: 5’-cacccgtcttcgatacatcaaacagc-3’; R: 5’-aaacgctgtttgatgtatcgaagac-3’

sgRNA-control; F: 5’-caccgttcctaagatttttaagact-3’; R: 5’-aaacagtcttaaaaatcttaggaac-3’.

### qPCR

qPCR was performed on 10 ng of DNA extracted from monolayer grown cells as described above. DNA and relevant primers were added to PowerUp SYBR Green Master Mix (Applied Biosystems) and real-time PCR performed using 7500 Fast Real-Time PCR System, using sybr green reagent. Primer sequences: HPV16 E2 F 5’ ATGGAGACTCTTTGCCAACG-3’ HPV16 E2 R 5’-TCATATAGACATAAATCCAG-3’; HPV16 E6 F 5’-TTGAACCGAAACCGGTTAGT-3’ HPV16 E6 R 5’-GCATAAATCCCGAAAAGCAA-3’

### SYBR green qRT-PCR

RNA was isolated using the SV Total RNA Isolation System (Promega) following the manufacturer’s instructions. Two micrograms of RNA were reverse transcribed into cDNA using the High Capacity Reverse Transcription Kit (Applied Biosystems). cDNA and relevant primers were added to PowerUp SYBR Green Master Mix (Applied Biosystems) and real-time PCR performed using 7500 Fast Real-Time PCR System. Primer sequences: SAMHD1 F 5’-ctggaactccatcccgactac-3’ SAMHD1 R 5’-agtaatgcgcctgtgatttcat-3’ GAPDH F 5’-ggagcgagatccctccaaaat-3’ GAPDH R 5’-ggctgttgtcatacttctcatgg-3’

### Protein analysis

1×10^6^ cells were lysed in 50 ul NP40 lysis buffer (0.5% Nonidet P-40, 50 mM Tris, pH 7.8, 150 mM NaCl) supplemented with protease inhibitor (Roche Molecular Biochemicals) and phosphatase inhibitor cocktail (Sigma). The cell and lysis buffer mixture was incubated on ice for 20 min, centrifuged for 20 min at 184,000 rfc at 4°C, and supernatant was collected. Protein levels were determined utilizing the Bio-rad protein estimation assay (Bio-rad). Equal amounts of protein were boiled in 2x Laemmli sample buffer (Bio-rad). Samples were then loaded into a Novex 4-12% gradient Tris-glycine gel (Invitrogen), run at 100V for approximately 2 hours, and then transferred onto nitrocellulose membranes (Bio-rad) at 30V overnight using the wet blot method. Membranes were blocked in Odyssey blocking buffer (diluted 1:1 with PBS) at room temperature for 6 h and probed with relevant antibody diluted in Odyssey blocking buffer overnight at 4°C. Membranes were then washed with PBS supplemented with 0.1% Tween (PBS-Tween) before probing with corresponding Odyssey secondary antibody (goat anti-mouse IRdye800cw or goat anti-rabbit IRdye680cw) diluted 1:10000 for 1h at 4°C. Membranes were washed in PBS-Tween before infrared scanning using the Odyssey CLx Li-Cor imaging system. The following antibodies were used for western blot analysis: GAPDH (1:10000, Santa Cruz sc-47724), SAMHD1 (1:1000, Cell Signalling Technology), V5 (1:500, Abcam).

### Organotypic raft culture

NOKs, NOKs+HPV16 and HTK+HPV16 cells were differentiated via organotypic raft culture as described previously (48, 49). Briefly, cells were seeded onto type 1 collagen matrices containing J2 3T3 fibroblast feeder cells. Cells were then grown to confluency on top of the collagen matrices, which were then lifted onto wire grids and cultured in cell culture dishes at the air-liquid interface, with media replacement on alternate days. Following 13 days of culture, rafted samples were fixed with formaldehyde (4% v/v) and embedded in paraffin blocks. Multiple 4μm sections were cut from each sample. Sections were stained with hematoxylin and eosin (H&E) and others prepared for immunofluorescent staining as described previously. Fixing and embedding services in support of the research project were generated by the VCU Massey Cancer Center Cancer Mouse Model Shared Resource, supported, in part, with funding from NIH-NCI Cancer Center Support Grant P30 CA016059.

### Immunofluorescence

Antibodies used and relevant dilutions are as follows: Involucrin (1/1000, Abcam), SAMHD1 (1:1000, Cell Signaling Technology), BrdU (1:200, Cell Signaling Technology), Cyclin E (1:1000, Sigma Aldrich), V5 (1/500, Abcam). Immune complexes were visualized using Alexa 488-or Alexa 595-labeled anti-species specific antibody conjugates (Molecular Probes). Cellular DNA was stained with 4’,6-diamidino-2-phenylindole (DAPI, Santa Cruz sc-3598). Fluorescent in situ hybridization (FISH) staining for HPV16 genomes was performed using DIG-labeled HPV16 genomes, as described previously (50, 51). Microscopy and subsequent quantification was performed at the VCU Microscopy Facility, supported, in part, by funding from NIH-NCI cancer center grant P30 CA16059. Immunofluorescence was quantified, using a Vectra^®^ Polaris™ automated imaging system, whereby whole stained sections were scanned computationally and the intensity calculated compared to a negative background control (secondary antibody only) and a positive localization control (DAPI). The same imaging parameters were used for each slide. For each sample, two sections from three individual rafts were scanned, to generate average values. Immunofluorescence was observed using a LSM 710 Laser Scanning Microscope and ZEN 2011 software (Carl Zeiss). Images were assembled in Adobe Photoshop CS 6.0.

### Southern Blot

Total cellular DNA was extracted using a phenol chloroform method and 5 micrograms digested with either *Sph*I or *Hind*III, to linearise the HPV16 genome or leave episomes intact, respectively. All digests included *Dpn*I to ensure that all input DNA was digested and not represented as replicating viral DNA. Digested DNA was separated by electrophoresis of a 0.8% agarose gel, transferred to a nitrocellulose membrane and probed with radiolabeled (32-P) HPV16 genome. This was then visualized by exposure to film for 24 or 72 hours.

### Statistics

Standard error was calculated from three independent experiments and significance determined using a student’s t-test.

## Acknowledgements

This work was supported by VCU Philips Institute for Oral Health Research and the National Cancer Institute Designated Massey Cancer Center grant P30 CA016059.

